# Zebrafish sleep displays distinct sub-states

**DOI:** 10.1101/2025.08.28.672887

**Authors:** Richa Tripathi, Grigorios Oikonomou, David A. Prober, Geoffrey J. Goodhill

## Abstract

Sleep is an essential and evolutionarily-conserved behavior. While mammals and several other species have been shown to exhibit well-defined sleep sub-states, some of which have been ascribed specific functions, it remains unclear to what extent such differentiation exists across the animal kingdom. Here we show, using long-term behavioral data combined with Hidden Markov Modeling, that larval zebrafish display distinct deep and light sleep sub-states. Although both states occur primarily at night, fish respond differently to sleep deprivation and arousing stimuli depending on which sleep sub-state they are in. Moreover, the proportions of deep and light sleep are selectively altered by genetic and pharmacological manipulations of melatonin, serotonin, and norepinephrine signaling, offering new insights into how these neuromodulators shape sleep architecture. These results support zebrafish as a tractable model for dissecting the regulation and function of sleep sub-states. More broadly, they demonstrate that structured, multi-state sleep is a conserved feature of vertebrate behavior.

## 1 Introduction

Sleep is regulated by a complex interplay of homeostatic and circadian mechanisms [1]. Based on the behavioral definition of sleep as a rapidly reversible and homeostatically regulated state of behavioral quiescence that is associated with an increased arousal threshold, sleep has been observed in all studied animals. However, despite being an evolutionarily conserved behavior [2, 3], the mechanisms that underlie sleep remain unclear [4, 5]. Mammalian sleep is not a uniform state but consists of distinct substates, which are linked to unique physiological processes spanning cellular, circuit, and systems levels [6–9]. Sleep sub-states have been identified in birds [10], reptiles [11–13], cephalopods [14], jumping spiders [15] and flies [16–18], however whether they also exist in other species remains unclear [2, 5, 19]. Zebrafish are a diurnal vertebrate offering key advantages for studying sleep, including at the larval stage their compatibility with whole-brain neuronal imaging and large-scale behavioral assays, and ease of genetic and pharmacological manipulation [20–22]. However the use of non-invasive electrophysiology such as EEG, the gold standard for identifying sleep sub-states in humans, is challenging in larval zebrafish. Recent work based on neural calcium imaging [23] has suggested the presence of sleep sub-states. However the use of highly constrained environments and visible wavelengths of light for fluorescence excitation leave open the possibility of confounds including stress responses, suppression of sleep by visible light, and homeostatic rebound effects.

An alternative approach for identifying sleep sub-states is based just on behavior, in particular overall levels of motor activity. Sleep is a quiescent state that should not be disturbed during measurement, thus to preserve its natural dynamics sleep is ideally studied non-invasively in freely behaving animals [24–26]. This approach not only allows for the characterization of normal sleep and its sub-states under standard conditions, but also leverages the advantages of zebrafish for high-throughput investigation of how genetic and pharmacological perturbations affect sleep [27, 28].

Here we exploit large-scale, long-term behavioral data of larval zebrafish combined with Hidden Markov Models (HMMs) to reveal distinct sleep sub-states. Using the Bayesian Information Criterion (BIC) we show that larval zebrafish locomotor activity is robustly optimally classified into 4 states, of which two are sleep sub-states (occurring primarily at night) and two are wake sub-states (occurring primarily during the day). Arousal assays and sleep-rebound experiments demonstrate that the two sleep sub-states match the characteristics of deep and light sleep states identified in other species. We then use this approach to reveal how the balance between deep and light sleep is altered by a range of genetic, pharmacological and environmental manipulations that affect sleep. Together, this work introduces a robust framework for identifying and analyzing sleep sub-states in larval zebrafish, and provides new insights into how neuromodulators shape sleep architecture in a diurnal vertebrate.

## 2 Results

Unless stated otherwise all analyses were based on the following experimental paradigm [25]. Larval zebrafish, raised under 14:10-hour light-dark (LD) conditions, were placed into 96-well plates at 4 days post-fertilization (dpf), and allowed to acclimate to their new environment overnight. Locomotor activity was then monitored for two days and nights, using infrared light and an infrared camera at 30 Hz, starting when white lights turn on at 9 a.m. at 5 dpf (Fig. 1a). Locomotor activity was based on pixel changes between consecutive frames (see Methods) and was quantified as seconds of locomotor activity per minute. We first analyzed wild-type (WT) fish, which show higher levels of locomotor activity during the day, when white lights are on, than at night, when white lights are off (Fig. 1b-d). Mean locomotor activity had a heavy-tailed distribution with a peak at zero (Fig. 1e).

**Figure 1:**
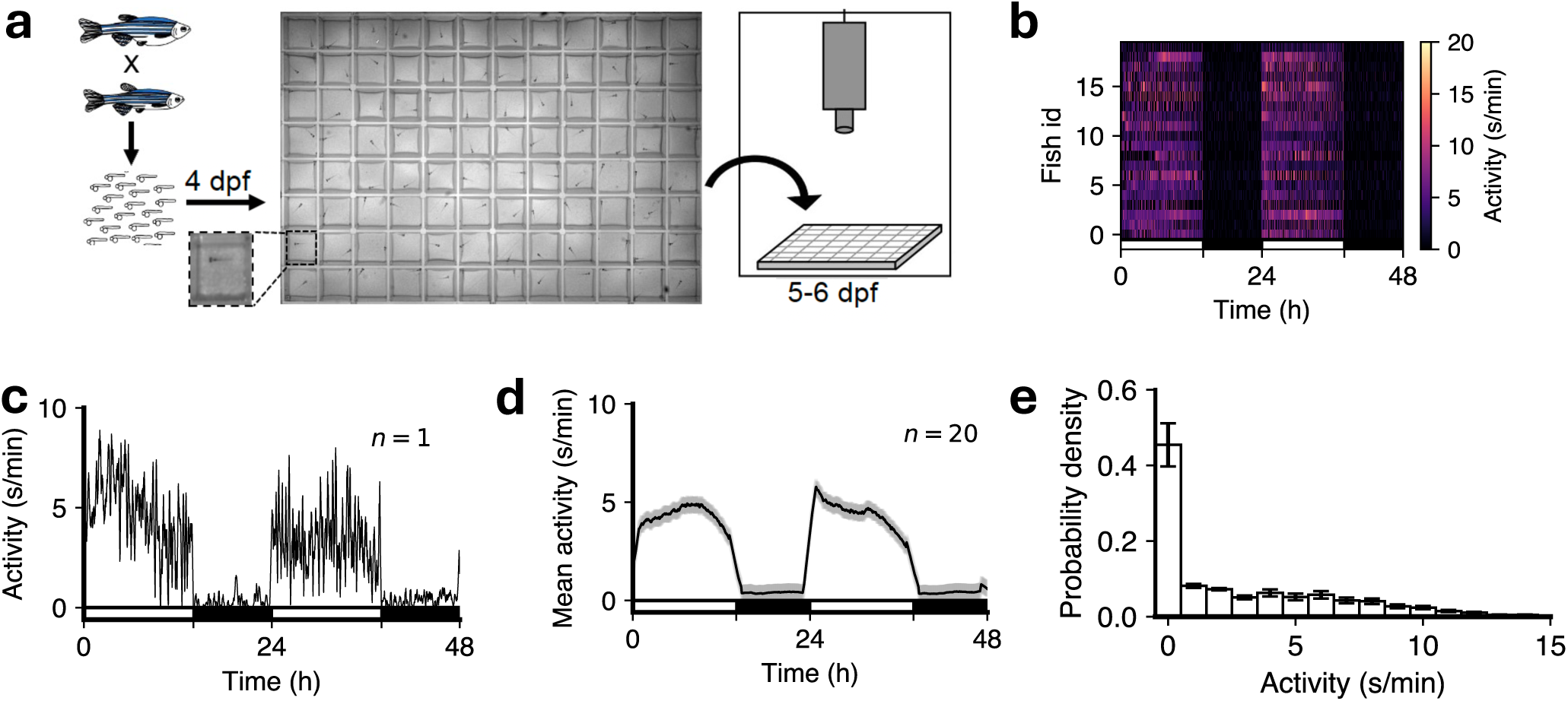
Monitoring larval zebrafish activity over 48-h day-night cycles. (**a**) Overview of experimental assay. (**b**) Heat-map of locomotor activity starting at 5 dpf (n = 21 fish). (**c**) Locomotor activity for one representative fish. (**d**) Mean activity for all fish, with sem shown in grey. (**e**) Distribution of locomotor activity for all fish (mean ± sem). In (**b-d**) the white and black horizontal bars depict the lights-on and lights-off periods on both the days, respectively.

### 2.1 Larval zebrafish display 2 sleep sub-states and 2 wake sub-states

To model these data we used HMMs. An HMM assumes that a system is progressing over time through a sequence of internal states which are not directly observed (“hidden”). Each state is associated with a distribution (here assumed to be Poisson) over observation probabilities, where here observations are activity in s/min. At each 1-min time step the system can stay in the same state or move to any of the other states with particular probabilities. The Markov assumption, which makes model fitting tractable, is that these transition probabilities, and thus the probability of the next state, depend only on the current state, and not on any longer history of the system. Using well-established algorithms it is straightforward to fit the sequence of states and transition probabilities that is most likely to have generated a particular sequence of observations. While the number of underlying states is assumed when fitting the model, an optimal number of states can be determined using the Bayesian Information Criterion (BIC). This trades off the better data fit that inevitably arises from models with more parameters against the increased complexity of the model, in order to find the simplest model that fits the data well.

However HMM fitting is not a deterministic process, and may get stuck in local minima. To help ameliorate this issue, in all cases reported here we ran the fitting algorithm 1000 times on each fish, and selected the model with the maximum log-likelihood of the fitted data (Fig. S1a). To determine if the amount of data available for each fish (48 h × 1 min = 2880 observations) was sufficient to reliably recover model parameters we first fitted 4-state HMMs to fish data. From these fitted parameters we then generated surrogate observation sequences of lengths ranging from 500 to 3500 minutes, and fitted HMMs to these. By 2880 observations the difference between the fitted and ground-truth parameters was small (Fig. S1b), giving us confidence that the length of our data was sufficient for reliable model fitting.

To determine the optimal number of states we fit HMMs containing 2-6 states to each fish and used BIC to determine which number of states was optimal. Interestingly there was some variation in the optimal number between fish (Fig. 2a). To determine if any of this variability was due to model fitting getting stuck in local minima we generated 100 state transition sequences and associated observations from a 4 HMM fit to a fish, refit those observations with HMMs with 2-6 states, and again used BIC to determine the optimal number of states. In 94% of cases the answer was again 4, indicating that the variation in optimal number of states between fish represents genuine biological variability rather than variability in model fitting (Fig. S1c). Consistent with this notion, there was considerable variation in sleep/wake behaviors among different animals (Fig. 1b). However despite this variability the modal optimal number of states across fish was 4, and the mean BIC value averaged over fish also had an optimum at 4 (Fig. 2a). For consistent subsequent analysis of HMM parameters we used 4-state HMM fits for all fish unless otherwise stated.

**Figure 2:**
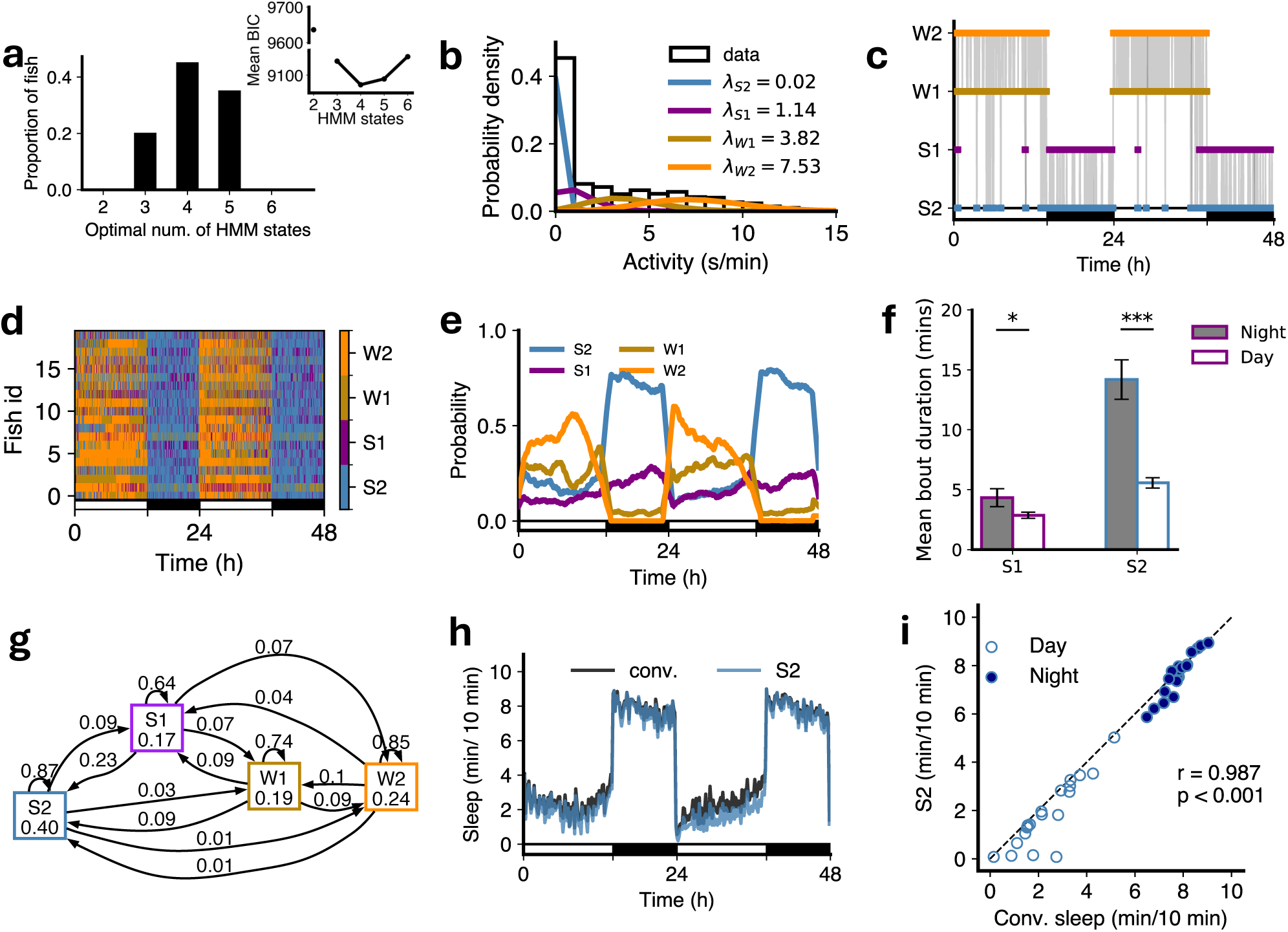
An HMM with 4 states best fits the data for WT fish. (**a**) The distribution of optimal number of states. Inset: Mean BIC as a function of number of HMM states. (**b**) The four means (*λ*s; units s/min) characterize the Poisson fits to fish locomotor activity for each state. (**c**) The most likely state sequence inferred from the data for a single example fish, based on parameters of a four state HMM fit. (**d**) State sequences for 21 fish over 48 hours at 5-6 dpf. (**e**) Variation of mean posterior probabilities of each state as a function of time, shown as a moving window average. (**f**) Both mean S1 and mean S2 bout durations were longer at night than during the day (paired t-test, *p*<*0.05, ***p*<*0.001, with Benjamini-Hochberg multiple comparisons correction). (**g**) The transition probabilities averaged across all fishes individual 4-state HMM fit. (**h**) Sleep amounts obtained based on the conventional definition of sleep (1 minute of no locomotor activity, black) compared to the most-likely state sequence from 4 HMM fit. (**i**) Comparison of HMM-defined S2 sleep with conventionally-defined sleep during day and night. Filled and empty circles represent night and day sleep amounts, respectively (one dot per fish). In (**c**), (**d**), (**e**) and (**h**), the white and black horizontal bars depict the lights on and lights off periods, respectively. r and p-value are for Pearson correlation for the fit to the line y=x.

For these 4 states the fitted means of the Poisson observation distributions (i.e. the mean amount of locomotor activity in each state, which we refer to as *λ*) were roughly 0, 1, 4 and 8 s/min (Fig. 2b). Following the terminology of [17] we refer to these states as S2, S1, W1 and W2 respectively. S2 and S1 were primarily observed at night and so we refer to them as sleep states, with wake states W1 and W2 observed primarily during the day (Fig. 2c-e, Fig. S2). The mean length of S1 and S2 states was also longer at night than during the day (Fig. 2f). The HMM transition diagram for the average fit over all fish is shown in Fig. 2g (discussed in next section).

The complete set of *λ* values for all fish is shown in Supplementary Tables 1-3. Supplementary Table 1 includes fish for which 4 states were optimal. Supplementary Table 2 includes fish for which 3 states were optimal, and also shows the *λ* values for the 4-state fit. In the 3-state fit these fish generally had only one high-activity (wake) state. Supplementary Table 3 includes fish for which 5 states were optimal, and also shows the *λ* values for the 4-state fit. In the 5-state fit these fish generally had an additional low-activity (sleep) state.

Sleep in larval zebrafish has been defined as any period of immobility longer than 1 min, because this is associated with an increase in arousal threshold [25, 29]. Comparing the occupancy of the S2 state with this conventional definition produced a good match (Fig. 2h,i). Note that fitting HMM states is not equivalent to simply defining states via locomotor activity thresholds. Consider for instance a 1 min bin with no activity. The relative probabilities that this was produced by Poisson distributions with the means for S2, S1, W1 and W2 found for the WT fish (0.02, 1.14, 3.82 and 7.53 s/min respectively) are 90.1%, 7.9%, 1.9% and 0.03% respectively. The model determines the most likely state assignment based on the complete sequence of state assignments, taking into account the tendency to persist in the same state between subsequent bins. Thus a bin with no activity could be classified as either S1 or S2 (and very occasionally W1). Across this set of fish, the fraction of bins with activity 0 s/min assigned states S2 and S1 were 0.84 and 0.16 respectively, and the fractions of bins with activity 1 s/min assigned as S2 and S1 were 0.09 and 0.82 respectively (with the remainder assigned to W1 or W2). This explains why the points in Fig. 2i lie slightly below the y = x line: the activity-0 bins S2 loses to S1 are not fully compensated for by the activity-1 bins S2 gains from S1, and thus the total S2 is slightly less than conventional sleep, i.e. the sum of all activity-0 bins.

The difference between the HMM model and defining states by simple activity thresholds is further emphasized by examining the distributions of activity assigned to each state (Fig. S3). For example there is considerable overlap between the distributions for S1 and W1 states, and the state assignment of a particular activity level depends on the broader context in which that activity level occurs. For instance, bins with activity of 2 s/min are assigned overall almost equally to S1 and W1 states. However at night they are assigned 84% of the time to S1 and only 15% of the time to W1, but during the day the assignment is 33% to S1 and 48% to W1 (Fig. S3g). Thus, the HMM captures contextual structure in the data that cannot be recovered by fixed activity thresholds.

For comparison we also examined the biological consequences of fitting these data with a 2-state HMM (Fig. S4), despite its much worse log-likelihood value. Unsurprisingly, the fitted *λ*s were intermediate between S1 and S2 for the ‘sleep’ state and intermediate between W1 and W2 for the ‘wake’ state. Because of the relatively high *λ* for the sleep state, the amount of time spent in this state was a poor match to conventionally-defined sleep. Thus, a simple 2-state model was both statistically suboptimal and biologically less informative compared to a 4-state model.

### 2.2 S1 is a light sleep state and S2 is a deep sleep state

Mammalian sleep consists of REM and NREM periods, and NREM sleep can be subdivided into distinct states characterized by different sleep depth. Deep NREM sleep has several behavioral properties that distinguish it from light NREM sleep. First, transitions to wake states are more likely to occur from light than deep sleep [17]. Second, although normally deep and light sleep states alternate, deep sleep is more prevalent under conditions of high homeostatic sleep pressure, such as the beginning of the night and during rebound sleep following sleep deprivation [30]. Third, animals in deep sleep are less responsive to an arousing stimulus than animals in light sleep [31]. We tested each of these properties for zebrafish to determine whether S2 represents a deep sleep state and S1 a light sleep state.

First, the HMM transition diagram (Fig. 2g) showed higher probabilities of transitioning from S1 to W1 (0.07) or W2 (0.07) than from S2 (0.03 and 0.01, respectively). Similarly, probabilities of transitioning from W1 to S1 (0.09) or S2 (0.09) were higher than probabilities of transitioning from W2 (0.04 and 0.01, respectively). Second, we conducted a sleep deprivation experiment where we exposed fish to daytime illumination for the first six hours of the second night of the experiment (Fig. 3a), which has been shown to strongly suppress sleep [32], and then turned the lights off. During the remaining 4 hours of the night there was increased sleep (i.e. rebound sleep) compared to the prior night of unperturbed sleep. We then fitted 4-state HMMs to determine how this affected S1 and S2 occupancies and transitions (Fig. S5a). Following lights off on the second night there was a large increase in S2 sleep at the expense of S1 sleep as compared to the first night (Fig. 3b,c). This suggests that the S2 state may play a crucial role in the homeostatic response to sleep deprivation in zebrafish.

**Figure 3:**
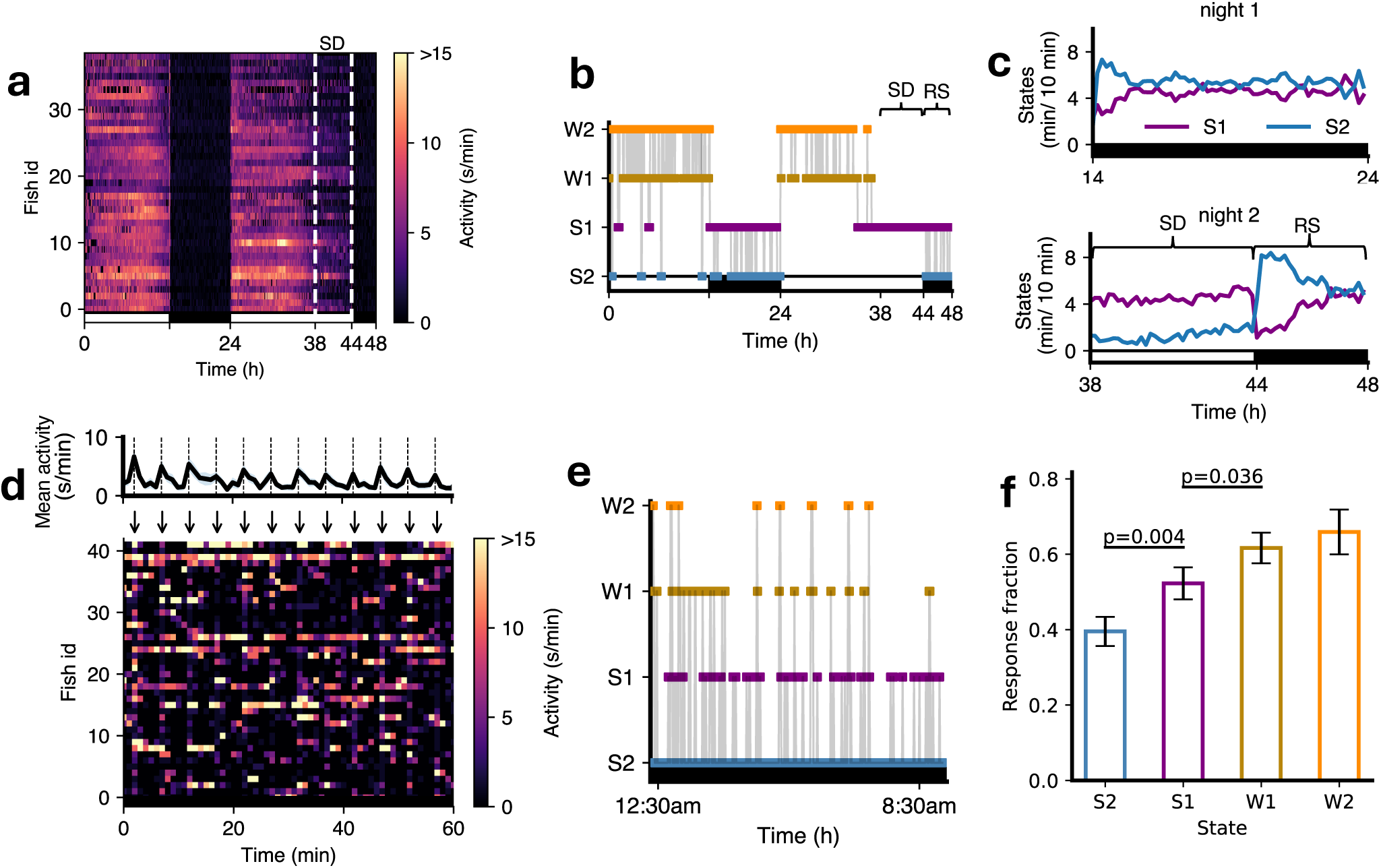
Fish in S1 vs S2 states show different responses to arousing stimuli and sleep deprivation. (**a-c**) Sleep deprivation experiment. (**a**) Fish were subjected to sleep deprivation (SD) on night 2, when lights were kept on for the first six hours at night (shown within dashed white lines), and then turned off for the last four hours of the night. (**b**) State sequence for an example fish over 48 hours (RS, rebound sleep). (**c**) Comparison of S2 and S1 during night 1 and night 2. (**d-f**) Arousal experiment. (**d**) An example 60 minutes of data with stimuli every 5 minutes (arrows) shown as a color map, with mean and standard error of mean locomotor activity over fish shown in upper plot. (**e**) A state sequence for an example fish over the night. In (**d**) and (**e**), the black horizontal bars indicate that lights were off. (**f**) Fish showed a higher proportion of responses (baseline-corrected) to stimuli when in the S1 state than S2 state, and when in the W1 state compared to the S1 state (t-tests with Benjamini-Hochberg multiple comparisons correction).

Third, we conducted an experiment where we exposed fish to a mechano-acoustic stimulus every five minutes during the night (Fig. 3d). We then fitted a 4-state HMM (Fig. 3e, S5b) to classify the sleep state just prior to each stimulus and determined the proportion of fish that showed any activity in the first two seconds after the stimulus, corrected for basal locomotor activity levels (see Methods). The proportion of fish that responded to the stimulus was significantly lower for S2 compared to S1, and for S1 compared to W1 (Fig. 3f). Thus, according to all three criteria, S2 represents a deeper sleep state than S1.

The latter experiment also allowed us to test whether HMM state assignments could differentiate arousability even for bins with the same absolute activity level. For this we exploited the overlap between the activity distributions assigned to S1 and W1 states. In particular, for this experiment we focused on bins preceding stimuli with activity 5 s/min, 40% of which were classified as S1 and 60% as W1. Remarkably, despite identical activity levels, fish were significantly less likely to respond to the stimulus when the state was classified as S1 compared to W1 (Fig. S5c). This further demonstrates the ability of the HMM to extract biologically meaningful underlying states in a way that exceeds the predictive power of simple thresholding.

In the remainder of this work we used the HMM approach to gain insight into how disruption of key mechanisms that regulate sleep affect sleep architecture, in particular states S1 and S2. Consistent with previous work using conventional sleep measures we found no significant differences between +/+ vs +/- fish for a given mutation (data not shown), so we focused our analyses on +/- vs -/- mutant comparisons.

### 2.3 Melatonin deficiency is associated with less deep sleep and more light sleep

Previous studies showed that melatonin acts downstream of the circadian clock to mediate circadian regulation of sleep in zebrafish [33]. Melatonin is synthesized in the pineal gland at night due to circadian expression of *aanat2*, which catalyzes the rate-limiting step of melatonin synthesis. Thus, melatonin-deficient *aanat2* mutant zebrafish can be used to study sleep architecture in the absence of circadian regulation of sleep. We first analyzed sleep architecture under standard 14:10 hour light-dark conditions, in which *aanat2^−/−^* fish were more active (Fig. 4a-c) and slept less at night (Fig. 4d) than their *aanat2*^+^*^/−^* siblings [33].

**Figure 4:**
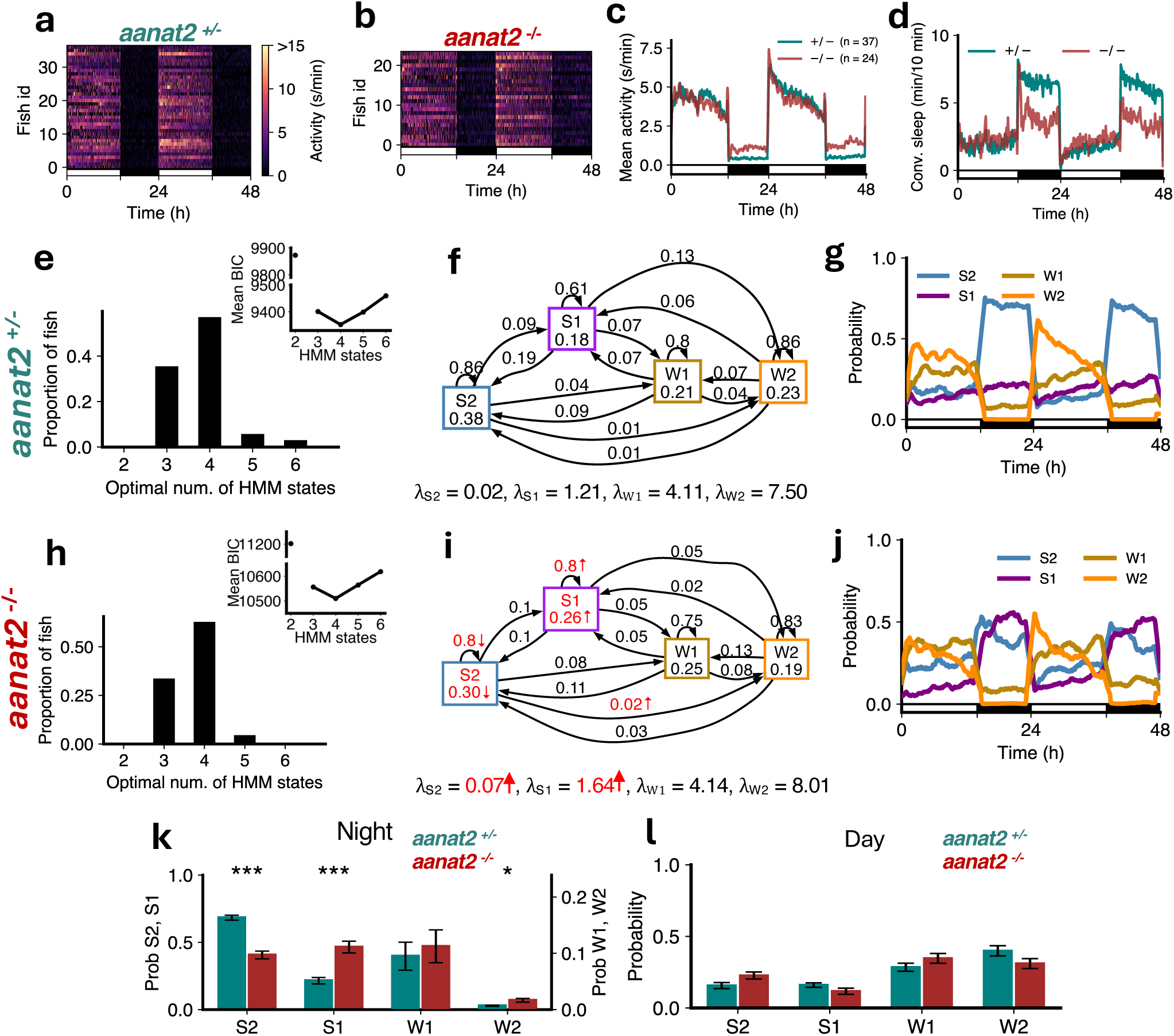
Melatonin-deficient *aanat2* mutants show loss of S2 and gain of S1. (**a-d**) *aanat2^−/−^* fish showed higher activity and lower sleep at night than *aanat2*^+^*^/−^* fish. (**e,h**) Both genotypes were best fit by 4 HMM states. (**f,i**) *aanat2^−/−^* fish had several significant differences in fitted HMM parameters (p *<* 0.05 indicated in red; bootstrap sampling with permutation tests; units of *λ*s: s/min). (**g,j**) *aanat2*^+^*^/−^* and *aanat2^−/−^* fish showed strikingly differing patterns of sleep during the night. (**k**) At night *aanat2^−/−^* fish had significantly less S2 but significantly more S1. (**l**) During the day there were no significant differences in genotypes between the times spent in each state. Asterisks indicate significant differences (t-tests, with Benjamini-Hochberg multiple comparisons correction; ***p*<*0.001, **p*<*0.01, *p*<*0.05).

For both *aanat2*^+^*^/−^* and *aanat2^−/−^* fish the optimal number of HMM states was 4 (Fig. 4e,h). Similar to WT fish (Fig. 2), sleep at night in *aanat2*^+^*^/−^* fish was dominated by S2 (Fig. 4g). However in *aanat2^−/−^* fish the amount of S2 was substantially reduced during the night (Fig. 4g, j, k, Fig. S6). Surprisingly the amount of S1 increased (Fig. 4g, j, k). The proportions of W1 and W2 were relatively unaffected, and there were no changes in any state probabilities during the day (Fig. 4l), as expected since little or no melatonin is produced during the day [33]. For *aanat2^−/−^* fish there was a significant increase in *λ_S_*_2_ (i.e. the mean locomotor activity while in the S2 state), and a significant decrease in the probability of occupying and remaining in S2 (Fig. 4f, i). There was also a significant increase in *λ_S_*_1_ and the probabilities of occupying and remaining in S1 for *aanat2^−/−^* fish (Fig. 4f, i). The reduction in S2 is consistent with conventional sleep analysis that *aanat2^−/−^* fish sleep less at night, but the model suggests this is mostly offset by an increase in S1 rather than wake states.

### 2.4 Free-running *aanat2* mutant fish show that light is required for state W2, and that melatonin is required for the increase in S2 that normally occurs at night

In order to analyze circadian regulation of behavior, it is necessary to first entrain animals in light:dark cycles, and then transfer them to constant dark, known as ‘free-running’ conditions. Under these conditions, entrained molecular and behavioral circadian oscillations persist for several days, allowing sleep to be studied without the masking effect of light on behavior [34]. Using this approach, circadian regulation of sleep is abolished in *aanat2* mutant zebrafish [33].

For both *aanat2*^+^*^/−^* and *aanat2^−/−^* fish overall activity levels were reduced in constant dark (Fig. 5a-c), as expected due to the loss of light-induced arousal. However during the subjective night, *aanat2^−/−^* fish had more activity and less sleep than *aanat2*^+^*^/−^* fish (Fig. 5c, d). Interestingly, the optimal number of states for the HMM fit of *aanat2*^+^*^/−^* was now only 3 (Fig. 5e,f). The lowest-activity state *λ* was similar to *λ_S_*_2_ for the unperturbed WT case (Fig. 2b) and this state showed an increase in probability during subjective night (Fig. 5g); we therefore assigned this state as S2. The highest-activity state had a *λ* similar to W1 for the unperturbed WT case (Fig. 2b); this state also showed an increase in probability during subjective day and we therefore labeled this state as W. The intermediate-activity state showed no obvious variation with subjective day and night, however its *λ* was similar to *λ_S_*_1_ for the unperturbed WT case (Fig. 2b) and so we assigned this state as S1. Thus constant dark has the effect of reducing the number of distinct sleep/wake states by effectively eliminating W2. This implies that the arousing effect of light is required for W2 states.

**Figure 5:**
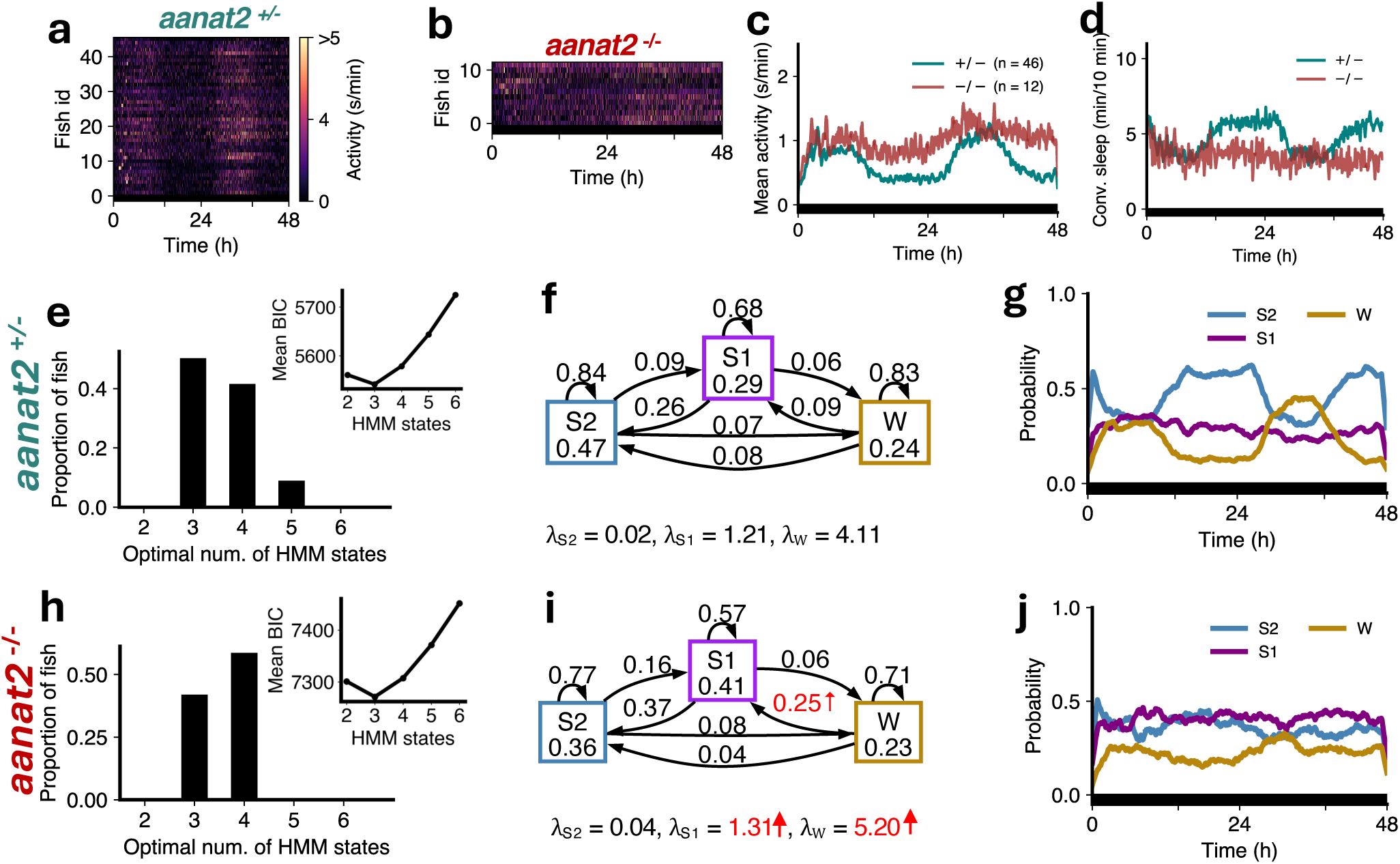
Circadian regulation of sleep at night is due to melatonin-dependent S2. (**a-d**) Overall activity was reduced under constant darkness, but *aanat2^−/−^* fish showed higher activity and lower sleep, and lacked circadian oscillations of sleep, at night compared to *aanat2*^+^*^/−^* fish. (**e,h**) Both genotypes had minimum BIC at 3 HMM states. (**f,i**) *aanat2^−/−^* fish showed significant changes in fitted HMM parameters (p *<* 0.05 in red; bootstrap sampling with permutation tests; units of *λ*s: s/min). (**g,j**) *aanat2*^+^*^/−^* fish maintained circadian variation in S2 and W states, but *aanat2^−/−^* lost circadian variation in all states.

For *aanat2^−/−^* fish the optimal number of states as defined by the mean BIC over fish was also 3 (Fig. 5h). There was a slightly higher number of individual fish with an optimum at 4 states (7 fish) rather than 3 states (5 fish), but for ease of comparison with *aanat2^−/^*^+^ fish we analyzed 3-state fits. These fish had significantly higher *λ_S_*_1_ and *λ_W_* than *aanat2*^+^*^/−^* Fig. 5f, i), consistent with generally higher activity levels (Fig. 5c). They also showed a significantly higher transition probability from W to S1. However none of these states showed any circadian variation Fig. 5j), demonstrating that circadian regulation of S2 is abolished by loss of melatonin. Based on these results, we conclude that melatonin mediates circadian regulation of sleep by increasing the S2 state at night.

### 2.5 Fish that lack serotonin in the raphe nuclei exhibit less deep sleep and altered sleep structure

The serotonergic raphe nuclei play an important role in homeostatic regulation of sleep in both zebrafish and mice [32]. *tph2* mutant zebrafish, whose raphe nuclei do not synthesize serotonin, are more active than sibling controls (Fig. 6a-c), sleep less (Fig. 6d) [32], and show reduced rebound sleep following sleep deprivation [32]. Thus, *tph2* mutants can be used to examine sleep architecture in the context of reduced homeostatic sleep pressure. We found that the optimal number of states that fit the data was 4 for both *tph2*^+^*^/−^* controls (Fig. 6e) and their *tph2^−/−^* siblings (Fig. 6h). However *tph2^−/−^* fish had larger *λ* values for all 4 states, with significant increases for S1, W1 and W2 (Fig. 6f,i). Additionally, *tph2* mutants had a lower probability of remaining in S2, lower occupancy of S2, and a higher probability of transitioning from S2 to S1 (Fig. 6f,i). *tph2* mutants also had higher probability of remaining in S1, and higher occupancies of S1 and W1 states (Fig. 6f,i). The S2 amount decreased and W1 amount increased during the day (Fig. 6g,j,l). At night, there was a significant reduction in the proportion of S2, and an increase in S1, in *tph2^−/−^* fish (Fig. 6g,j,k). Thus, while conventional analysis indicates a reduction in sleep in *tph2* mutants, HMM analysis instead indicates a shift in balance between deep and light sleep. This observation suggests that serotonin plays a key role in promoting deep sleep, consistent with previous findings of lighter sleep in *tph2* mutants [32], and with the observation that increased homeostatic sleep pressure results in an increase in deep sleep at the expense of light sleep (Fig. 3c).

**Figure 6:**
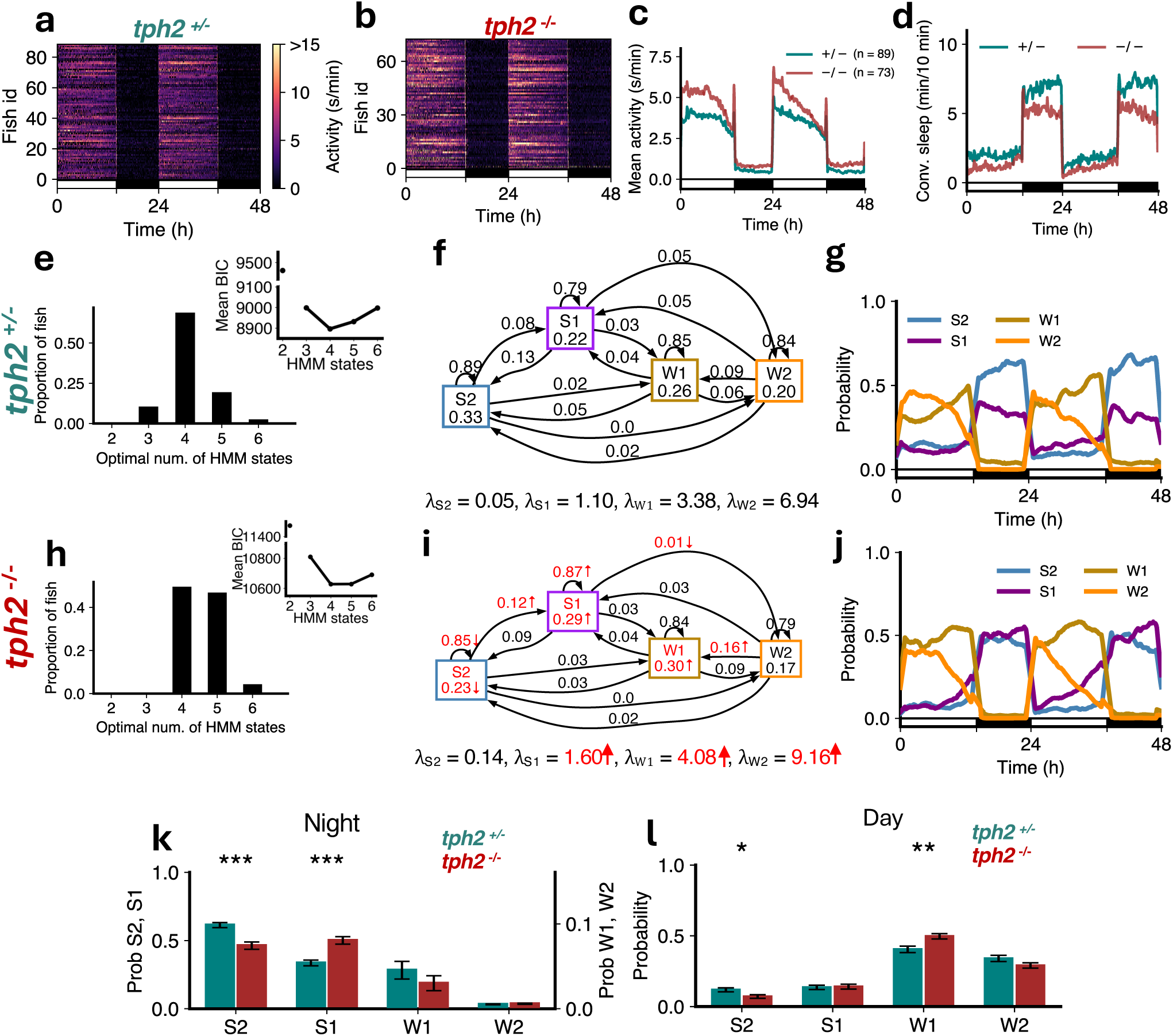
*tph2* mutants show less deep sleep. (**a-d**) *tph2^−/−^* fish showed higher activity and lower sleep than *tph2*^+^*^/−^* fish. (**e,h**) Both genotypes were best fit by 4 HMM states. (**f,i**) *tph2^−/−^* fish had several significant differences (p *<* 0.05 indicated in red; bootstrap sampling with permutation tests) in fitted HMM parameters (*λ* units: s/min). (**g,j**) *tph2*^+^*^/−^* and *tph2^−/−^* fish showed similar patterns of sleep states during the day but differing patterns at night. (**k**) At night *tph2^−/−^* fish had significantly less S2 but significantly more S1. (**l**) During the day there were no significant differences in genotypes between the times spent in each state. Asterisks indicate significant differences (t-tests, with Benjamini-Hochberg multiple comparisons correction; ***p*<*0.001, **p*<*0.01, *p*<*0.05).

### 2.6 Pharmacological activation of serotonin signaling results in more deep sleep

To further explore the role of serotonin in regulating sleep sub-states, we analyzed the behavior of fish treated with quipazine, a serotonin receptor agonist, compared to DMSO vehicle-treated controls. Quipazine treatment results in reduced locomotor activity (Fig. 7a-c) and increased sleep compared to controls (Fig. 7d) [32]. For both groups the optimal number of HMM states was 4 (Fig. 7e,h). All *λ* values were reduced (all significantly except *λ_S_*_1_), consistent with reduced overall activity. The amount of S2 was dramatically increased at night, with a concomitant loss of S1 (Fig. 7g,j,k), while there were no significant differences during the day (Fig. 7l). Consistent with this there were significant increases in the probabilities of staying in the S2 state and transitioning from S1 to S2, and a decrease in the probability of transitioning from S2 to S1 (Fig. 7f,i). These results are consistent with the *tph2* mutant data (Fig. 6), and with the hypotheses that serotonin promotes deep sleep but not light sleep, and that increased homeostatic sleep pressure promotes deep sleep at the expense of light sleep.

**Figure 7:**
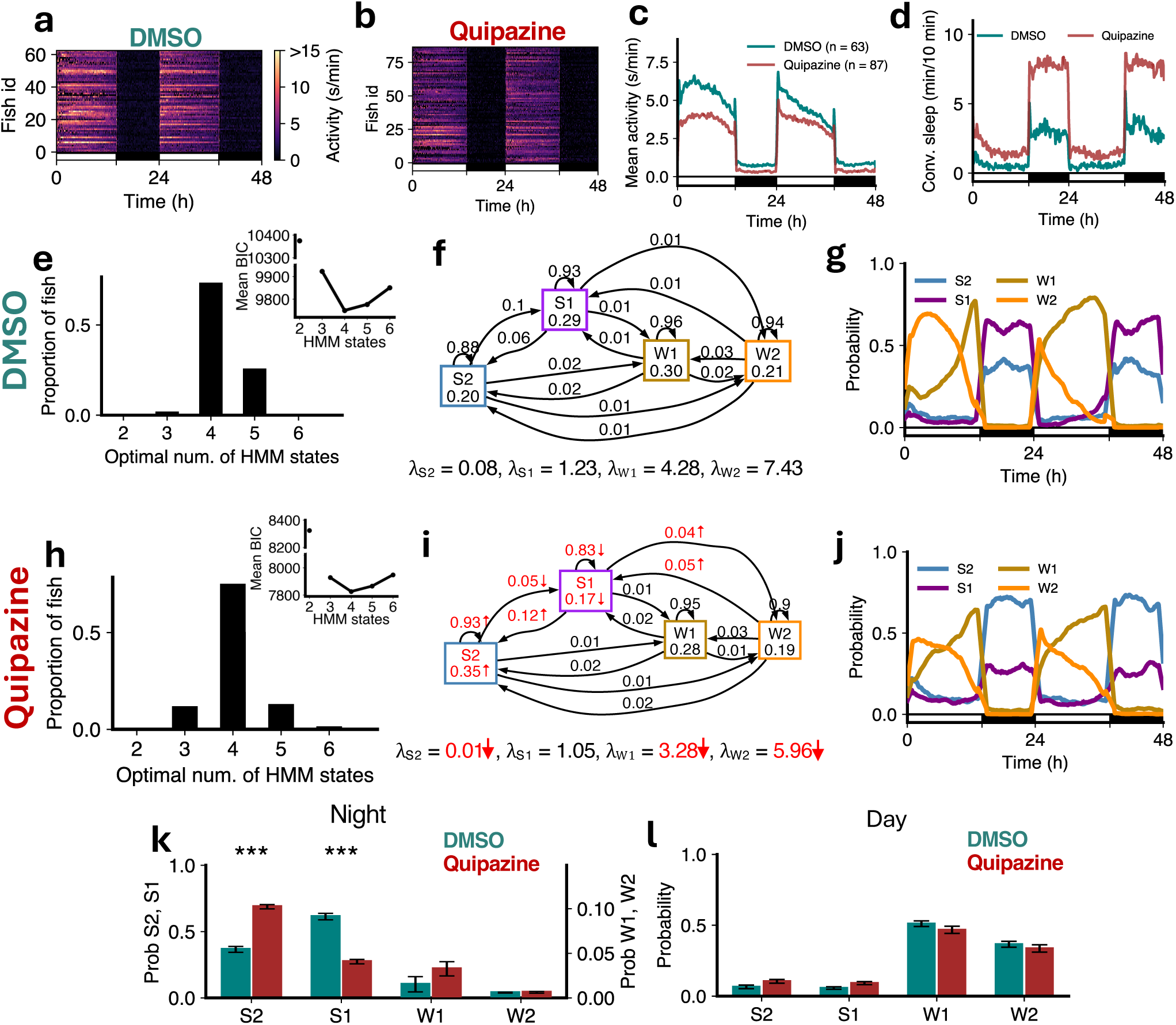
Quipazine treatment induces sleep through gain in deep sleep. (**a-d**) Quipazine treated fish showed lower activity and more sleep than DMSO controls. (**e,h**) Both genotypes were best fit by 4 HMM states. (**f,i**) Quipazine fish had several significant differences (p *<* 0.05 indicated in red; bootstrap sampling with permutation tests) in fitted HMM parameters (*λ* units: s/min). (**g,j**) Quipazine fish showed changes in sleep states during the night. (**k**) At night quipazine fish had significantly higher S2 but significantly lower S1. (**l**) During the day there were no significant differences in groups between the times spent in each state. Asterisks indicate significant differences (t-tests, with Benjamini-Hochberg multiple comparisons correction; ***p*<*0.001).

### 2.7 Noradrenaline-deficient *dbh* mutant fish sleep more due to increased deep sleep

Noradrenaline is a major wake-promoting neurotransmitter that helps maintain arousal and suppresses sleep across vertebrate species including zebrafish [35]. *dbh^−/−^* fish, which lack norepinephrine, were less active (Fig. 8a-c) and slept more (Fig. 8d) than sibling *dbh*^+^*^/−^* controls [35]. Both mutants and controls were best fit by 4 HMM states (Fig. 8e,h). While there were no significant changes in the *λ*s for S2, S1 and W1, *λ_W_* _2_ increased significantly in *dbh^−/−^* fish. *dbh^−/−^* fish had a significant increase in the occupancy of S2, and a decrease in the probability of transitioning from S2 to S1 (Fig. 8f,i), leading to their altered proportions during both day and night (Fig. 8g,j,k,l). The probabilities of remaining in wake states dropped significantly (Fig. 8f,i), resulting in a significant reduction in the proportions of W1 and W2 during the day (Fig. 8l). Notably, W2 occupancy and the probability of transitioning from W2 to S2 and S1 increased significantly for *dbh^−/−^* fish (Fig. 8f,i). These results suggest that loss of noradrenaline results in increased deep sleep at the expense of light sleep.

**Figure 8:**
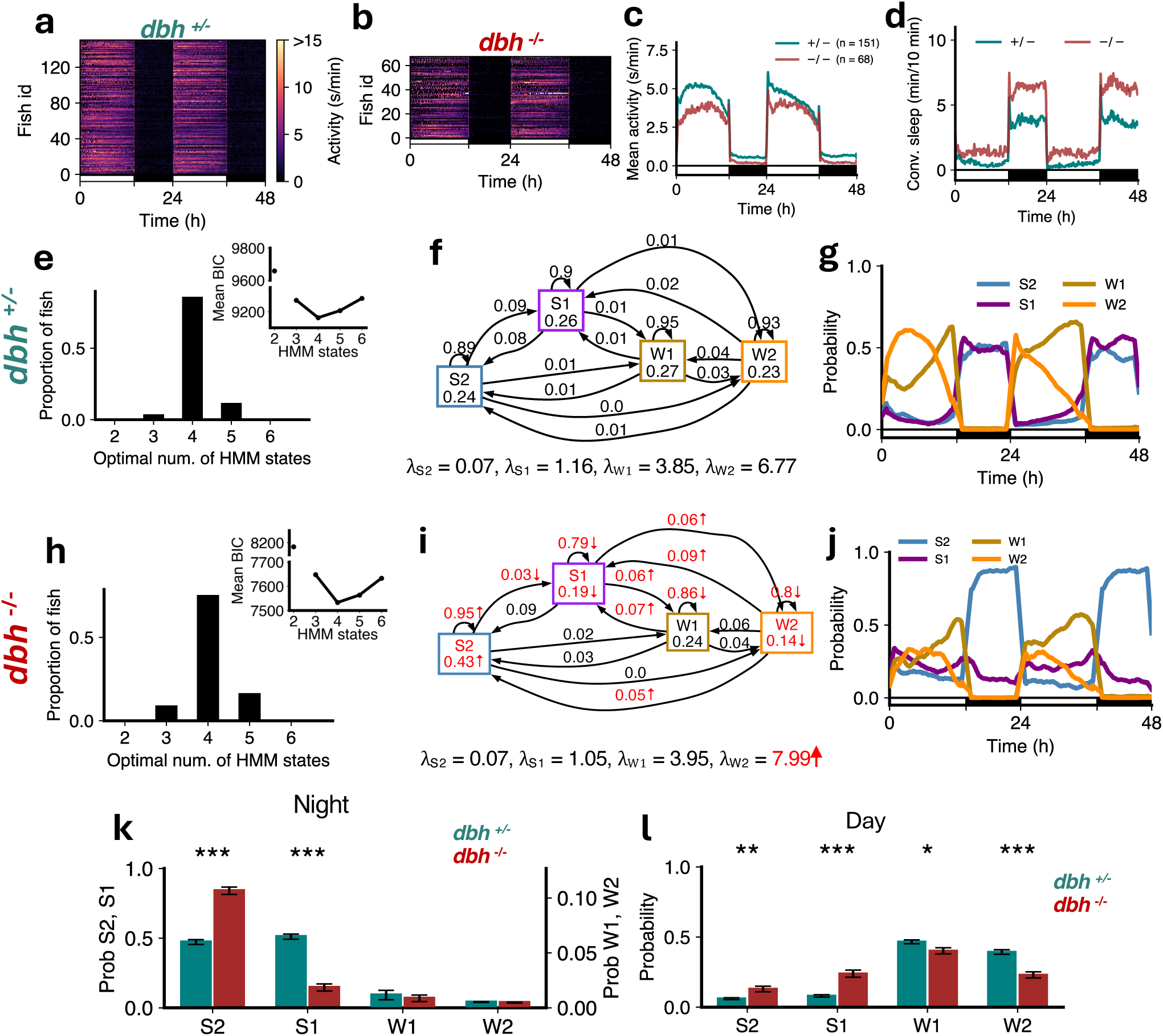
dbh mutants show more deep sleep and less wake occupancy. (**a-d**) *dbh^−/−^* fish showed lower overall activity and higher sleep than *dbh*^+^*^/−^* fish. (**e,h**) Both genotypes were best fit by 4 HMM states. (**f,i**) *dbh^−/−^* fish had several significant differences (p *<* 0.05 indicated in red, bootstrap sampling with permutation tests) in fitted HMM parameters (*λ* units: s/min). (**g,j**) *dbh*^+^*^/−^* and *dbh^−/−^* fish showed altered patterns of sleep during night and day and wake states during the day. (**k**) At night *dbh^−/−^* fish had significantly high S2 but significantly less S1. (**l**) During the day there were significant increases in S2 and S1, and a decrease in W2, in *dbh^−/−^* fish. Asterisks indicate significant differences (t-tests with Benjamini-Hochberg multiple comparisons correction; ***p*<*0.001, **p*<*0.01, *p*<*0.05)

### 2.8 Pharmacological inhibition of noradrenaline signaling phenocopies *dbh* mutant behavior

Prazosin is an alpha1-adrenergic receptor antagonist, which blocks the postsynaptic effects of noradrenaline alpha1 receptors, and thus phenocopies *dbh* mutant fish [35]. Prazosin-treated fish had lower levels of locomotor activity (Fig. 9a-c) and slept more at night than controls (Fig. 9d) [35]. In both cases the optimal number of HMM states was 4 (Fig. 9e,h). In the HMM transition diagram there was a significant increase in the occupancy of S2, decrease in occupancies of S1, W1 and W2, accompanied by increases in the probability of transitioning from W2 to lower activity states, and a decrease in the probability of remaining in S1, W1 and W2 (Fig. 9f,i). There were significant changes in the amounts of S1 and S2 both during night and day in the drug-treated fish (Fig. 9g,j,k,l), indicating that the S2 occupancy change was also driven by daytime changes. Consistent with this, during the day there was a large increase in the proportion of S2 and S1 and a decrease in the proportion of W1 and W2 (Fig. 9l). Consistent with the *dbh^−/−^* fish, prazosin-treated fish did not show significant differences in *λ*s except for the *W* 2 state (Fig. 9f,i). Since noradrenaline signaling normally promotes arousal during the day, this suggests that alpha1-adrenergic tone suppresses transitions into deep sleep during the active phase. These results support a model in which noradrenaline, via alpha1 receptors, promotes wakefulness by limiting inappropriate or premature entry into deep sleep when animals would otherwise be active. Compared with *dbh* mutant fish (Fig. 8k, l), prazosin-treated fish showed similar changes in states both during the night and day, consistent with the established role of noradrenaline signaling.

**Figure 9:**
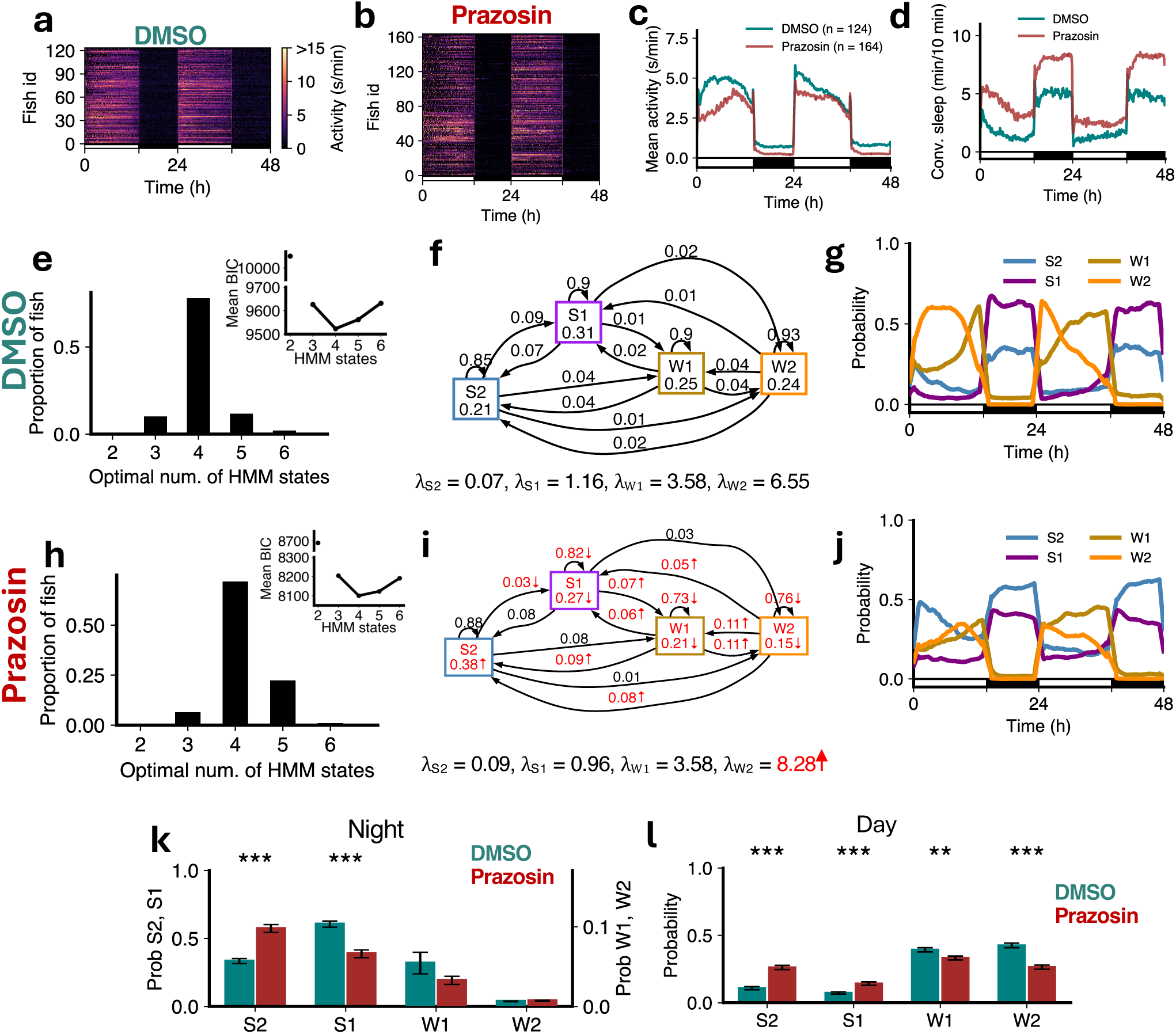
Prazosin treatment induces sleep through gain in deep sleep and loss of wake. (**a-d**) Prazosin-treated fish showed lower activity and higher sleep than DMSO controls. (**e,h**) Both groups were best fit by 4 HMM states. (**f,i**) Prazosin-treated fish had several significant differences (p *<* 0.05 indicated in red, bootstrap sampling with permutation tests) in fitted HMM parameters (*λ* units: s/min). (**g,j**) Prazosin fish showed an increase in S2 during the night and day, and reduced W2 during the day. (**k**) At night prazosin fish showed a significant increase in S2 and a significant decrease in S1. (**l**) During the day there were significant differences in groups between the times spent in each of S2, S1 and W2. Asterisks indicate significant differences (t-tests with Benjamini-Hochberg multiple comparisons correction; ***p*<*0.001, **p*<*0.01, *p*<*0.05).

## 3 Discussion

Although recent computational analyses of freely moving zebrafish have shed light into the temporal organization of behavior [36–40], none have specifically addressed sleep. Here, by applying HMMs to long-term locomotor recordings, we reveal that zebrafish sleep comprises two distinct sub-states. These states occur predominantly at night, display distinct timing, transition dynamics, arousability and rebound following sleep deprivation, and are differentially modulated by neuromodulatory systems. This richer characterization aligns with evidence from other species that sleep comprises multiple functional substates rather than a unitary condition. By establishing a robust computational framework for dissecting sleep architecture in larval zebrafish, this work paves the way for further insights into sleep regulation.

Traditional definitions of zebrafish sleep, based on quiescence thresholds, are primarily captured by S2, with *λ_S_*_2_ ≈ 0 s/min (Fig 1i). The S1 state is characterized by non-zero but much lower levels of activity than wake states (*λ_S_*_1_ ≈ 1 s/min, as compared to *λ_W_* _1_ ≈ 4 s/min and *λ_W_* _2_ ≈ 8 s/min). The statistically-derived value of *λ_S_*_1_ ≈ 1 s/min matches with an earlier analysis which heuristically specified a 3-state model where between 0 and 1 s/min of activity was defined as a ‘low-activity’ state and more than 1 s/min activity as a ‘high-activity’ state [25]. While S1 generally occurs more at night it shows the surprising property of being increased by manipulations which reduce total sleep defined conventionally in zebrafish (Figs 4,6). Indeed, changes in S1 often appear to compensate for changes in S2 (Figs 4,6,7,8). This follows naturally from the structure of the HMM model: reductions in S2 caused by a small increase in overall activity will mostly be captured by S1, since the activity increase required to be captured by the W states is larger. This raises the question of whether S1 is best regarded as a true sleep state (which is supported by our arousal threshold data (Fig 3f), or perhaps instead as a drowsy or quiet wake state. This could potentially be addressed by higher-resolution analysis of more subtle behaviors, such as eye, tail and fin movements, heart rate, respiration rate and postural control, though these experiments would be much lower throughput than the simple locomotor activity data we have analyzed here.

Ultimately electrophysiological recordings will be required to fully determine how S1 and S2 states compare with standard definitions of mammalian sleep states. Leung et al [23] recorded calcium activity in the dorsal pallium of head-restrained zebrafish larvae using 1-photon microscopy and reported states somewhat analogous to NREM and REM, the latter of which they termed propagating wave sleep. However, these states were only observed after prolonged sleep deprivation, perhaps because natural sleep was inhibited by the requirement to restrain the animals for imaging, or by the intense visible light required to excite GCaMP flourescence. Our data suggests that zebrafish S1 and S2 states are more analogous to light and deep stages of NREM sleep in mammals than REM and NREM respectively, although higher resolution behavioral data, and neuronal activity monitoring in a manner that does not require bright visible light, are needed to fully address this question.

HMMs have emerged as a powerful computational framework for analyzing sleep architecture across diverse species, leveraging their ability to capture the temporal dynamics and probabilistic transitions between distinct behavioral states. In human sleep research, HMMs have been extensively applied to automate sleep stage classification from polysomnographic recordings, utilizing EEG, EOG, and EMG signals to identify transitions between sleep stages while accounting for the inherent temporal structure of sleep architecture [41, 42]. HMMs have also been adapted for sleep/wake identification using human actigraphy data, expanding their utility to large-scale studies where traditional polysomnography is impractical [43]. In mice four-state Markov models have been used to characterize sleep-wakefulness dynamics along light/dark cycles [44]. In invertebrate models, probabilistic analyses including HMM-based approaches have revealed covert sleep-related biological processes in *Drosophila*, enabling quantification of sleep pressure and sleep depth parameters [17]. The versatility of HMMs in capturing both observable behavioral patterns and inferring hidden states makes them particularly well-suited for comparative sleep studies across phylogenetically diverse species, providing a unified analytical framework that can accommodate the varied temporal scales and behavioral manifestations of sleep across different animal models. In our work we used BIC to determine the optimal number of states and were careful to address the non-deterministic nature of HMM fitting, fitting the model multiple times to each fish and then choosing the one with the highest log-likelihood.

Our modeling approach also revealed individual variability in sleep state architecture. While four was always the modal optimal number states (except in free-running conditions when the W2 state was lost), in each condition some fish showed three, five or even six states. However two states was never the optimal solution, showing that a binary classification into unitary ‘sleep’ and ‘wake’ states is an incomplete characterization of zebrafish sleep-wake cycles. The variation in the number of optimal states that we observed between individual fish is not surprising given the documented individual differences in sleep/wake behaviors in fish [25, 26, 45] and mammals [46]. These observations suggest that zebrafish will be useful for future studies investigating interindividual variation in sleep phenotypes.

We also observed variability in the proportions of S1 and S2 between different control groups. Although for each experiment the mutant/drug case was performed at the same time as the controls, we have collectively analyzed data collected over many years. This long-term variability likely reflects a combination of factors including individual differences, subtle environmental fluctuations and gradual evolution of video-tracking technology. Consider the case of a small decrease d in the amount of conventionally-defined sleep between two experiments being the result of a small increment in overall activity. Time bins lost to S2 will be captured mostly by S1, so that the S2:S1 ratio is expected to change by approximately (1-d)/(1+d) ∼ 1-2d, i.e. a change twice as large as the change in conventionally-defined sleep. However, the directional shifts we observed in sleep states were consistent across multiple independent manipulations targeting distinct neuromodulatory pathways, each compared with its respective control group. This consistency suggests that the effects we report reflect robust changes in sleep architecture rather than sampling noise. Future work could reduce batch variability by running multiple manipulations in parallel with shared controls.

By examining how specific genetic and pharmacological perturbations alter the occupancy and transitions of sleep states, we demonstrate that distinct neuromodulators play complementary roles in shaping zebrafish sleep architecture. Melatonin acts downstream of the circadian clock to support deeper sleep at night, emphasizing the importance of circadian output for sleep depth as well as timing. Serotonin appears critical for promoting progression into and maintenance of deep sleep: loss of serotonin in *tph2* mutants reduces S2, while the serotonin agonist quipazine shifts sleep in the opposite direction. Loss of noradrenaline increases deep sleep and decreases maintenance of wakefulness during the day. In contrast, at night noradrenaline-deficient mutants exhibit increased deep sleep and reduced light sleep. Together, these findings show that monoaminergic neuromodulators do not simply gate sleep-wake transitions but actively shape the internal architecture of sleep, biasing the balance between lighter and deeper sleep sub-states with time of the day.

A limitation of our approach is that the HMM infers hidden states based solely on statistical regularities in the behavioral data, without guaranteeing that the inferred states correspond directly to biologically meaningful categories. However previous work has demonstrated that HMMs applied purely to human actigraphy data can accurately reproduce sleep states derived from simultaneous polysomnography [43]. Moving beyond sleep, there is compelling evidence that HMMs applied to mouse behavioral data can reveal hidden states corresponding to known physiological categories [47]. This work demonstrates the power of behaviorally-based approaches for revealing physiological states, especially when physiological measurements are hard to obtain directly.

Together, our results suggest that the ability to structure sleep into depth-defined sub-states is an evolutionarily conserved feature that does not depend on mammalian-specific brain structures. More broadly, our work highlights the importance of analyzing sleep microarchitecture to uncover how neuro- modulatory pathways shape not just when animals sleep, but how deeply they sleep. This provides a more nuanced view of sleep regulation that bridges behavior, circuits, and evolution.

## 4 Material and Methods

### 4.1 Zebrafish experiments

Zebrafish were raised and maintained at 28.5*^◦^*C under 14:10 hours light:dark conditions with white lights on from 9 a.m. to 11 p.m. At 4 dpf, individual zebrafish were placed in each well of a 96-well plate, and locomotor activity and sleep were monitored for 48 h from 5-6 dpf, starting with lights on at 9 a.m. at 5 dpf, using a videotracking system as previously described [48]. Behavior was monitored at 30 Hz and data was integrated into 1 minute bins for the HMM analyses. Details on fish husbandry, matings, and genotyping can be found in previous papers that presented these data sets [32, 33, 35]. Sleep deprivation and acoustic stimulation were performed as described [32]. In the sleep deprivation experiment, during the second night of behavioral recording, the white lights that normally turned off at 11 p.m. were kept on for 6 hours until 5 a.m., and then turned off for 4 hours until 9 a.m. In the acoustic stimulus experiment, a stimulus was delivered every 5 minutes from 12:30 a.m. to 9 a.m. in the dark while behavior was monitored (99 stimuli total).

### 4.2 HMM fitting

We used Poisson HMMs (PoissonHMM module of hmmlearn library in Python) to fit the fish activity, which was quantified by the number of active seconds in each 60 second bin. For BIC calculations, we fitted 1000 models with different random seeds for each fish for each of 2-6 states, and selected the model in each case with the maximum log-likelihood. For each fish, state occupancies were estimated using the dominant eigenvector of the transition probability matrix. Average state probabilities over time were smoothed with a window of 60 min.

For the sleep deprivation and arousal experiments only 4-state HMMs were fitted (again the best of 1000 fits). For the arousal experiment, since the data per fish spanned only a single night, we concatenated data from all the fish together and then fitted a single 4-state HMM. From the fitted model we inferred state assignments over time for each fish. To allow a fair comparison we excluded fish which did not occupy each of the 4 states at least once during the experiment. This left 24 of an initial 42 fish remaining. The 1 min time bins were aligned so that each stimulus occurred just after the end of a bin. The state assigned to this previous bin was taken as the state of the fish at the time of the stimulus. The fish was recorded as having responded to the stimulus if there was any movement in the 2 s following the stimulus. To correct this initial probability for baseline activity (i.e. the probability that a fish in a particular state would have moved in a 2 s window without the stimulus) we applied the same procedure but based on the 1 min time bin starting 2 min before the stimulus, and then subtracted this probability from that calculated initially.

### 4.3 Statistical tests

To test for significant differences between HMM parameters for fish in different groups we performed bootstrap sampling from the two groups and a permutation test to determine a p value (with p *<* 0.05 being taken as significant). This procedure was applied for all state occupancies, *λ*s and transition probabilities. A Bonferroni correction was used for multiple comparisons (a factor of 4 for state occupancies and *λ*s, and 16 for transition probabilities).

## 5 Acknowledgements

This work was supported by the NIH grants UF1 NS126562 to D.A.P and G.J.G, and R35 NS122172 to D.A.P. We thank Bruno van Swinderen and Maxwell Shafer for feedback on the manuscript.

---

**Supplementary Table 1.** 𝛌 values for fish with 4 states as optimal.

| | $\lambda_{S2}$ | $\lambda_{S1}$ | $\lambda_{W1}$ | $\lambda_{W2}$ |
| --- | --- | --- | --- | --- |
| fish 1 | 0.02 | 1.27 | 4.65 | 7.04 |
| fish 2 | 0.01 | 0.23 | 0.91 | 5.89 |
| fish 3 | 0.01 | 1.10 | 5.76 | 8.23 |
| fish 7 | 0.01 | 1.33 | 3.67 | 8.97 |
| fish 9 | 0.01 | 1.00 | 4.97 | 8.86 |
| fish 10 | 0.00 | 1.27 | 4.18 | 5.87 |
| fish 11 | 0.02 | 1.16 | 3.85 | 7.08 |
| fish 18 | 0.02 | 0.94 | 3.17 | 8.52 |
| fish 19 | 0.02 | 1.35 | 5.72 | 8.70 |

**Supplementary Table 2.** 𝛌 values for fish with 3 states as optimal.

|  |  |  |  |  |  |  |  |
| --- | --- | --- | --- | --- | --- | --- | --- |
| fish 4 (3 state) | 0.02 |  | 1.55 |  | 7.00 |  |  |
| fish 4 (4 state) | 0.02 |  | 1.29 |  | 5.50 |  | 8.65 |
| fish 15 (3 state) | 0.03 |  | 1.79 |  | 8.91 |  |  |
| fish 15 (4 state) | 0.03 |  | 1.26 |  | 2.23 |  | 9.06 |
| fish 17 (3 state) | 0.01 |  | 2.15 |  | 7.24 |  |  |
| fish 17 (4 state) | 0.00 |  | 1.21 |  | 4.01 |  | 7.91 |
| fish 20 (3 state) | 0.01 |  | 1.52 |  | 4.77 |  |  |
| fish 20 (4 state) | 0.00 |  | 1.06 |  | 3.41 |  | 5.87 |

**Supplementary Table 3.** 𝛌 values for fish with 5 states as optimal.

|  |  |  |  |  |  |
| --- | --- | --- | --- | --- | --- |
| fish 5 (5 state) | 0.04 | 1.11 | 1.16 | 4.62 | 6.79 |
| fish 5 (4 state) | 0.04 | 1.16 | 1.36 | 5.33 |  |
| fish 6 (5 state) | 0.04 | 0.09 | 1.28 | 3.71 | 5.12 |
| fish 6 (4 state) | 0.05 | 1.49 | 3.92 | 5.13 |  |
| fish 8 (5 state) | 0.01 | 0.11 | 1.53 | 3.14 | 5.30 |
| fish 8 (4 state) | 0.01 | 0.31 | 1.54 | 4.28 |  |
| fish 12 (5 state) | 0.03 | 0.40 | 1.23 | 4.08 | 7.69 |
| fish 12 (4 state) | 0.03 | 1.23 | 2.49 | 6.97 |  |
| fish 13 (5 state) | 0.02 | 0.18 | 2.11 | 3.45 | 7.62 |
| fish 13 (4 state) | 0.04 | 2.11 | 6.35 | 9.56 |  |
| fish 14 (5 state) | 0.01 | 0.02 | 1.29 | 3.42 | 7.73 |
| fish 14 (4 state) | 0.00 | 1.36 | 4.93 | 8.22 |  |
| fish 16 (5 state) | 0.03 | 0.70 | 1.54 | 4.62 | 10.45 |
| fish 16 (4 state) | 0.05 | 1.50 | 5.27 | 10.63 |  |

**Figure S1:**
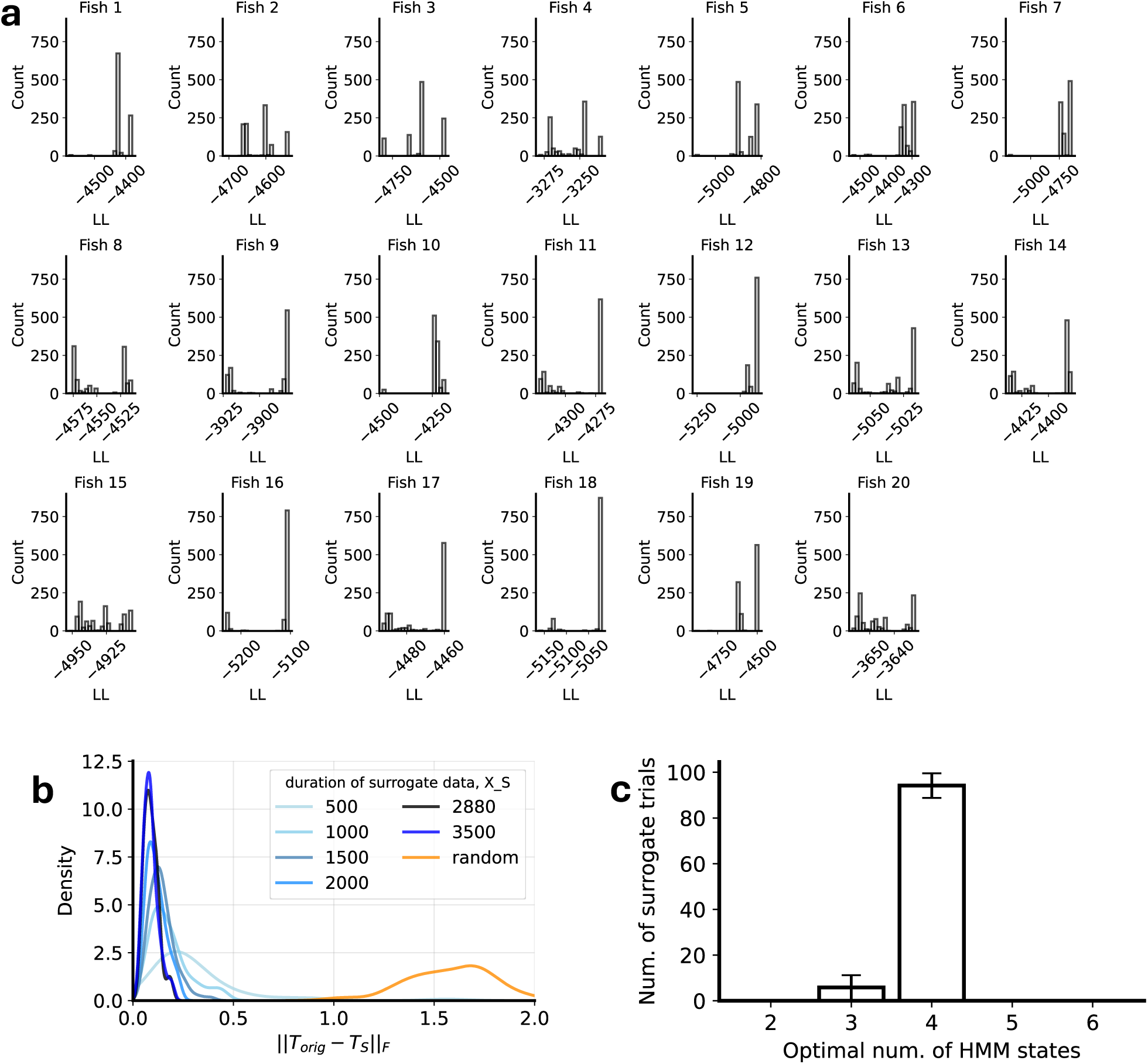
Robustness of HMM parameter fits. (**a**) Distribution of log-likelihood (LL) over 1000 model fits for each WT fish, depicting variability among different runs. (**b**) Distribution of difference (Frobenius norm) between the transition probability matrix *T_orig_* fitted on 48 h fish data and the transition probability matrix fitted on surrogate data generated from *T_orig_*, as a function of the amount of surrogate data provided. The distributions became narrower as longer durations of data were used for fitting, with little change after duration of 2880 minutes. For the random distribution case each row of the transition probability matrix was generated from a Dirichlet distribution and normalized to sum to one. (**c**) For surrogate data generated 100 times from a 4-HMM fit to a fish for which 4 states was optimal, the optimal number of fitted states was 4 in 94 cases. The error bars are standard error of mean over fish.

**Figure S2:**
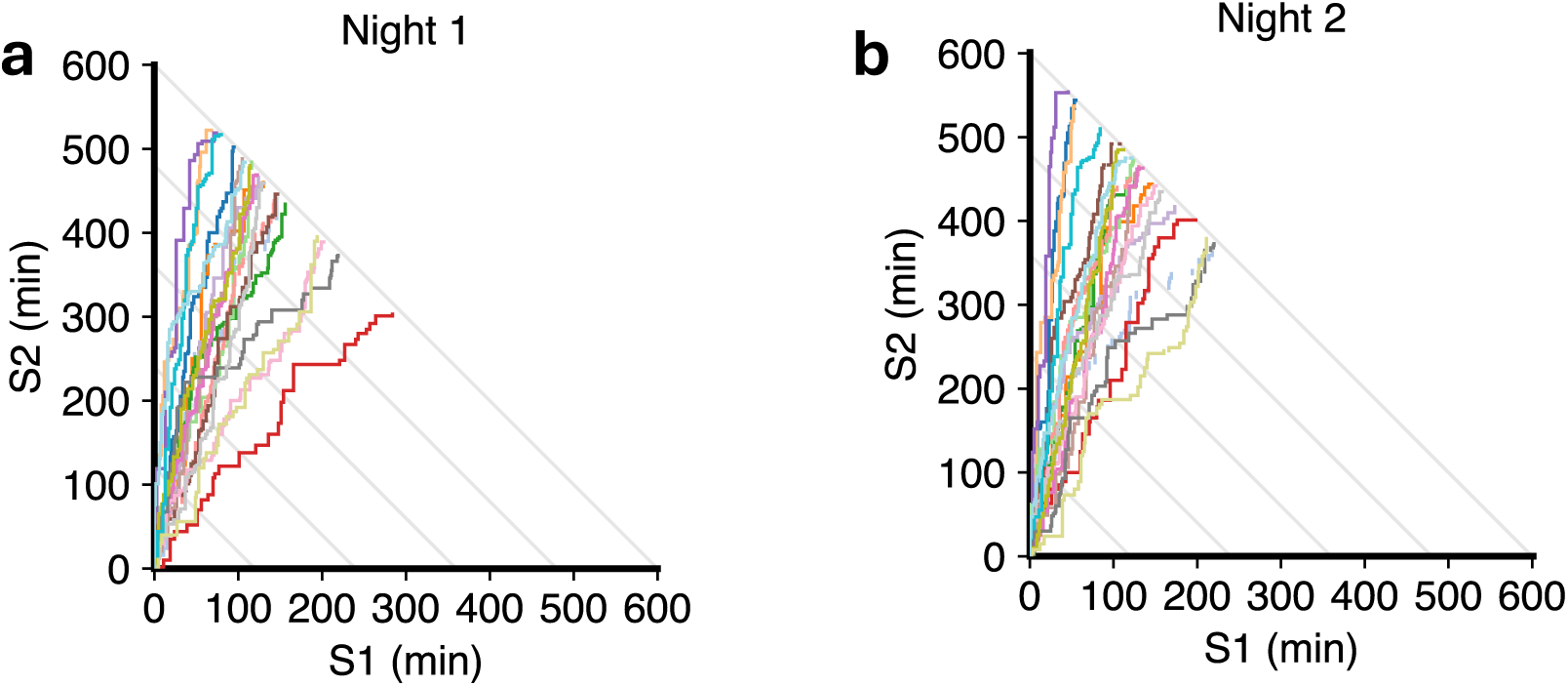
Complete S1 and S2 state sequences at night for all WT fish. In this representation night starts at the origin, and then we draw a horizontal line for each minute in state S1 and a vertical line for each minute in S2. Each color corresponds to a different fish (matched between nights). In the rare instances when a wake state occurs we leave a horizontal space. All lines thus finish at 600 total min (10 h). These diagrams provide a compact way of representing individual sleep-state sequences for all fish in the same plot.

**Figure S3:**
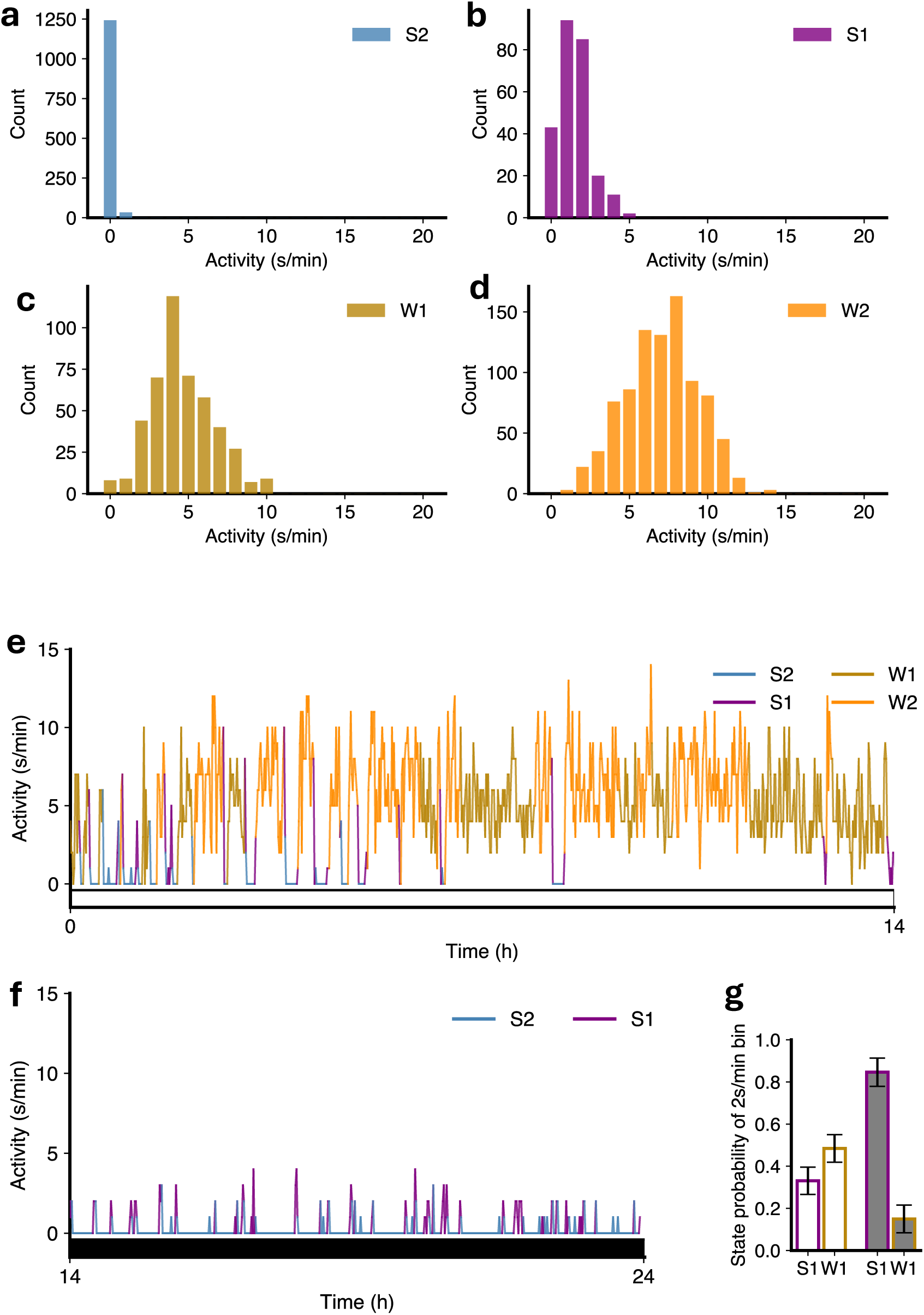
Relation of activity levels to state assignments and arousal probability. (**a-d**) Histograms for an example WT fish showing the distribution of activity levels assigned to each of the four states. (**e-f**) Activity levels of the same fish colored by the state assigned at each time during the first day (**e**) and night (**f**). (**g**) On average, bins with 2 s/min of activity are more likely to get assigned state W1 during the day (empty bars), but state S1 during the night (filled bars).

**Figure S4:**
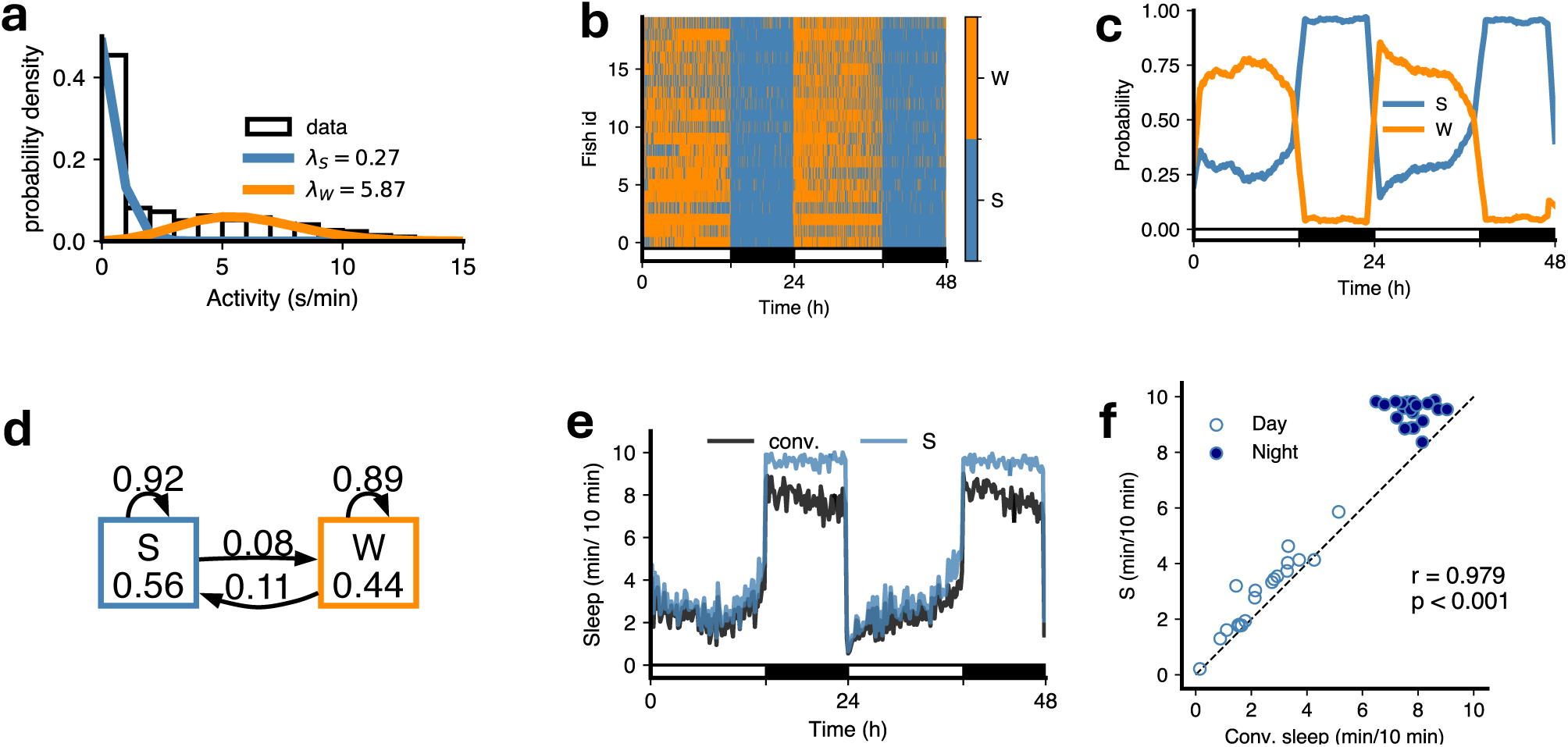
A 2-HMM fit to WT fish data. (**a**) *λ*s for Poisson distribution fit to activities in both states (S and W). (**b**) Most likely state sequences underlying fish activities. (**c**) State probabilities during day and night phases. (**d**) Transition diagram for the model. (**e**) A comparison of sleep amounts for the S state and sleep calculated conventionally. (**f**) HMM sleep versus conventional sleep for each fish during day and night. r and p-value are for Pearson correlation for the fit to the line y=x.

**Figure S5:**
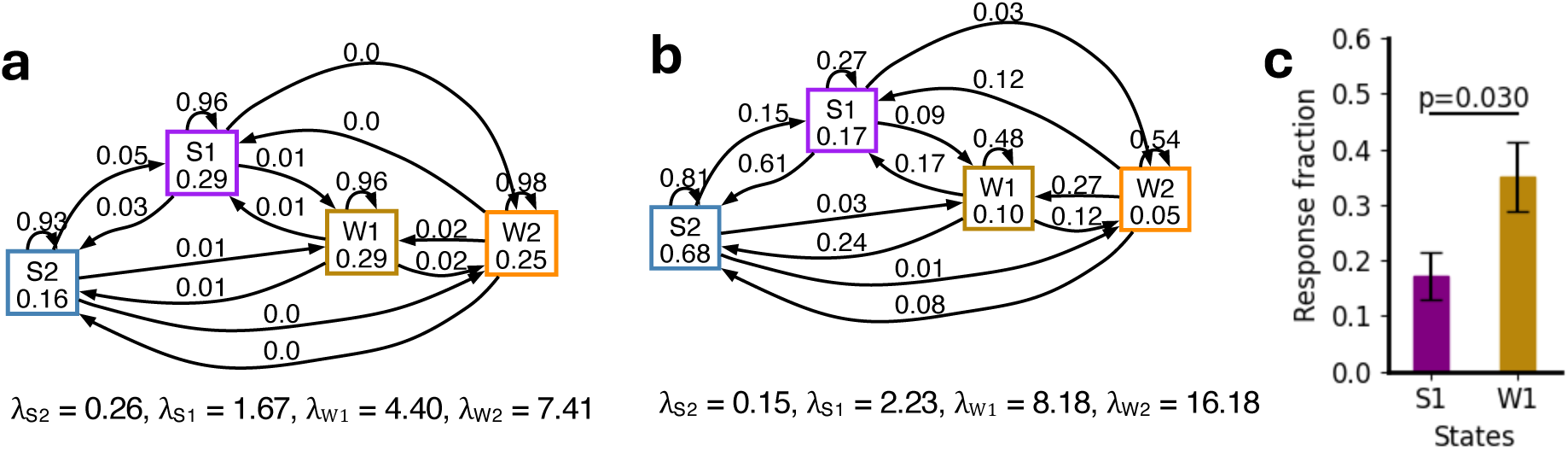
HMM parameters for sleep deprivation and arousal experiments. (**a**) Mean transition diagram and *λ* values for the HMM fit to sleep deprivation data. (**b**) Mean transition diagram and *λ* values for the HMM fit to arousal experiment data. (**c**) Response fractions from state S1 versus state W1, both corresponding to activity level of 5 s/min before the stimulus.

**Figure S6:**
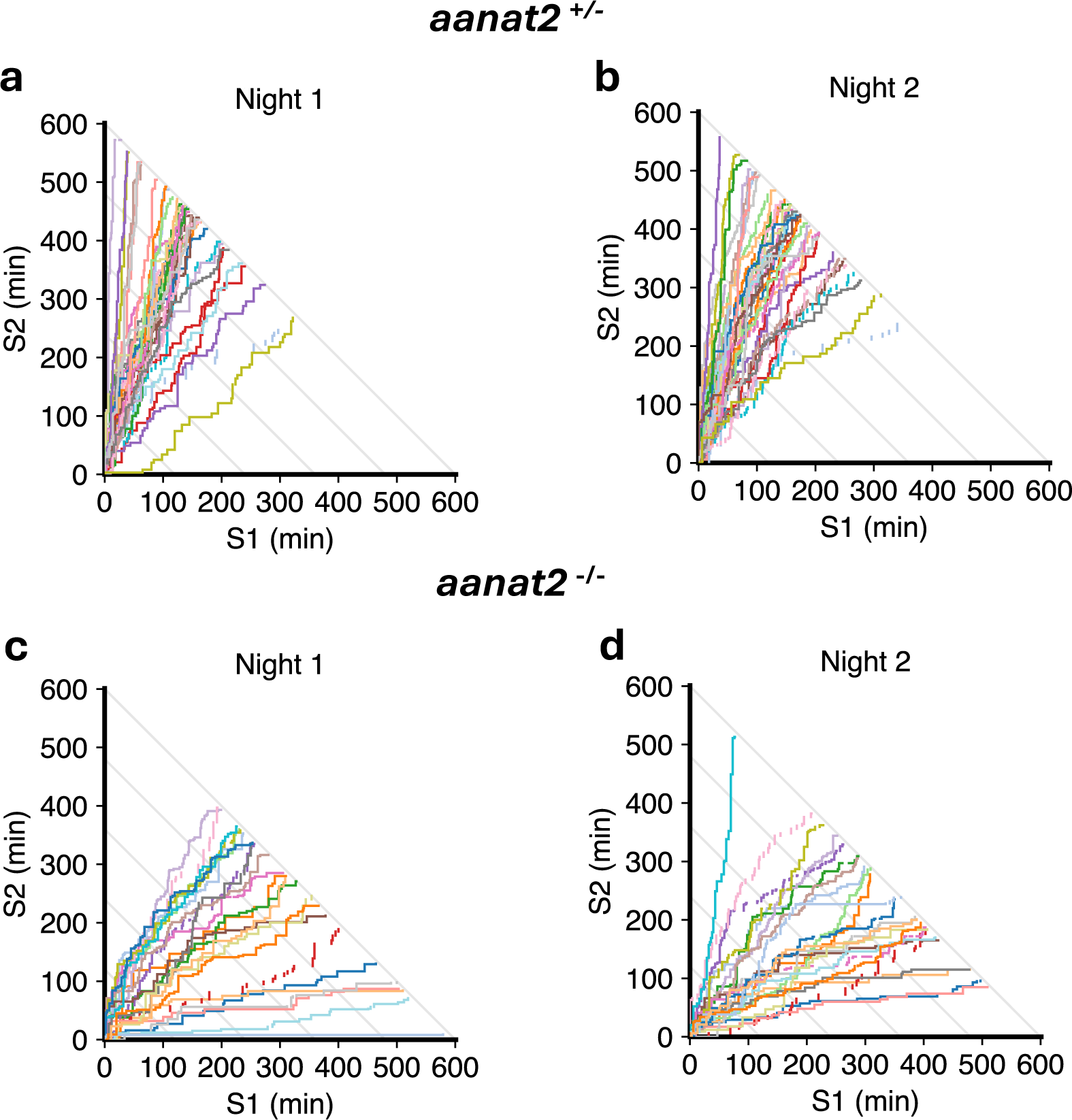
Comparison of S1 and S2 state sequences during night for *aanat2*^+^*^/−^* and *aanat2^−/−^* fish. (**a-b**) *aanat2*^+^*^/−^* fish. (**c-d**) *aanat2^−/−^* fish. Colors represent different fish. The increase in S1 relative to S2 in *aanat2^−/−^* fish is clearly apparent.

